# Conceptual models of entrainment, jet-lag, and seasonality

**DOI:** 10.1101/865592

**Authors:** Isao T. Tokuda, Christoph Schmal, Bharath Ananthasubramaniam, Hanspeter Herzel

## Abstract

Understanding entrainment of circadian rhythms is a central goal of chronobiology. Many factors, such as period, amplitude, *Zeitgeber* strength, and day-length, govern entrainment ranges and the phase of entrainment. Using global optimization, we derive conceptual models with just three free parameters (period, amplitude, relaxation rate) that reproduce known phenotypic features of vertebrate clocks: relatively small phase response curves (PRCs), fast re-entrainment after jet-lag, and seasonal variability to track light onset or offset. Since optimization found multiple sets of model parameters, we can study this model ensemble to gain insight into the underlying design principles. We find that amplitudes control the size of PRCs, that fast relaxation supports short jet-lag, and that specific periods allow reasonable seasonal phase shifts. *Arnold* onions of representative models visualize strong dependencies of entrainment on periods, relative *Zeitgeber* strength, and photoperiod.

## 1 INTRODUCTION

### 1.1 Entrainment and oscillator theory

The circadian clock can be regarded as a system of coupled oscillators, be it the neuronal network in the SCN (Hastings et al., 2018) or the ‘orchestra’ of body clocks (Dibner et al., 2010). Furthermore, the intrinsic clock is entrained by *Zeitgebers* such as light, temperature and feeding. The concept of interacting oscillators (Huygens, 1986; Van der Pol and Van der Mark, 1927; Kuramoto, 2012; Strogatz, 2004) can contribute to the understanding of entrainment (Winfree, 1980). The theory of periodically driven self-sustained oscillators is based on the concept of “*Arnold* tongues” (Pikovsky et al., 2003; Granada et al., 2009). *Arnold* tongues mark the ranges of periods and *Zeitgeber* strengths in which entrainment occurs (Abraham et al., 2010). The range of periods over which entrainment occurs is called the “range of entrainment” (Aschoff and Pohl, 1978). If seasonal variations are also considered, the entrainment regions have been termed “*Arnold* onions” (Schmal et al., 2015). Within these parameter regions, amplitudes and entrainment phases can vary drastically. Amplitude expansion due to periodic forcing is termed resonance (Duffing, 1918). Of central importance in chronobiology is the variability of the entrainment phase, since the coordination of the intrinsic clock phase with the environment provides evolutionary advantages (Aschoff, 1960; Ouyang et al., 1998; Dodd et al., 2005).

### 1.2 Phenomenological amplitude-phase models

After the discovery of transcriptional feedback loops (Hardin et al., 1990), many mathematical models focussed on gene-regulatory networks (Leloup and Goldbeter, 2003; Forger and Peskin, 2003; Becker-Weimann et al., 2004). However, most available data on phase response curves (PRCs) (Johnson, 1992), entrainment ranges (Aschoff and Pohl, 1978), and phases of entrainment (Rémi et al., 2010) are based on organismic data. Thus it seems reasonable, to study phenomenological models that are directly based on these empirical features. There is a long tradition of heuristic amplitude-phase models in chronobiology (Klotter, 1960; Wever, 1962; Pavlidis, 1973; Daan and Berde, 1978; Winfree, 1980; Kronauer et al., 1982).

Here, we examine the limits of the capability of such heuristic amplitude-phase models to reproduce fundamental properties of circadian entrainment. To this end, we combine the traditional amplitude-phase modeling approach with recent oscillator theory and global optimization to identify minimal models that can reproduce essential features of mammalian clocks: relatively small PRCs (Honma et al., 2003), fast re-entrainment after jet-lag (Yamazaki et al., 2000), and seasonal variability (Daan and Aschoff, 1975).

### 1.3 Empirical data as model constraints

In order to optimize phenomenological models, reasonable constraints based on experimental observations have to be formulated. In mammals, the strong coupling of SCN neurons constitutes a strong oscillator (Abraham et al., 2010; Granada et al., 2013) with quite small PRCs (Comas et al., 2006). Even bright light pulses of 6.7h duration can shift the clock by just a few hours (Khalsa et al., 2003). Consequently, we constrain our models to have small PRCs with just 1h advance and 1h delay. Interestingly, despite the robustness of the SCN rhythms, a surprisingly fast recovery from jet-lag is observed (Reddy et al., 2002; Vansteensel et al., 2003). Along the lines of a previous optimization study (Locke et al., 2008), we request that our models reduce the jet-lag to 50 % within 2 days. The third constraint refers to the well-known seasonal variability of circadian clocks (Pittendrigh and Daan, 1976b; Rémi et al., 2010). It has been reported that phase markers can lock to dusk or dawn for varying day-length. This implies that the associated phases change by 4h, if we switch from 16:8 LD conditions to 8:16 LD conditions. Thus, we test whether or not our optimized models allow such pronounced phase differences between 16:8 and 8:16 LD cycles.

## 2 METHODS

### 2.1 Optimization of amplitude-phase model

As a model of an autonomous circadian oscillator, we consider the following amplitude-phase model (Abraham et al., 2010):

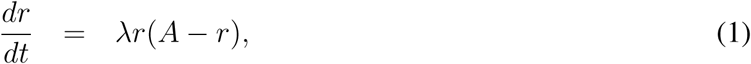

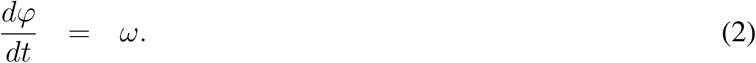

The system is described in polar coordinates of radius *r* and angle *ϕ*, having a limit cycle with amplitude *A* and angular frequency *ω*. Any perturbation away from the limit cycle will relax back with a relaxation rate *λ*. This oscillator model can be represented in Cartesian (*x, y*)–coordinates as

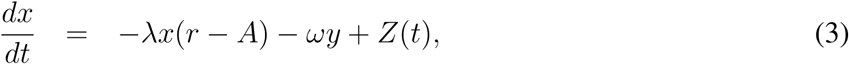

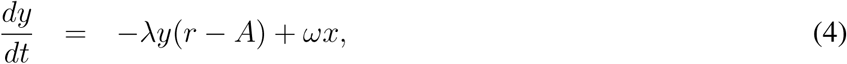

where 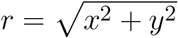. The oscillator receives a *Zeitgeber* signal

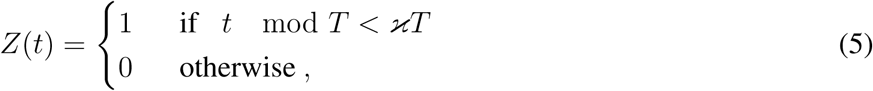

where *T* represents the period of the *Zeitgeber* signal and *κ* determines the photoperiod (*i.e.*, fraction of time during *T* hours when the lights are on). The amplitude–phase model provides one of the simplest mathematical frameworks to study limit cycle oscillations, which have been discussed in the context of circadian rhythms (Wever, 1962; Winfree, 1980; Kronauer et al., 1982).

The amplitude–phase model (1),(2) has three unknown parameters {*A, ω, λ*}. These parameters were optimized to satisfy the model constraints as described in 1.3. The parameter optimization is based on minimization of a cost function. The cost function takes a set of parameters as arguments, evaluates the model using those parameters, and then returns a “score” indicating goodness of fit. Scores may only be positive, where the closer a fit gets to the score of zero, the better the fit becomes. The cost function is defined as

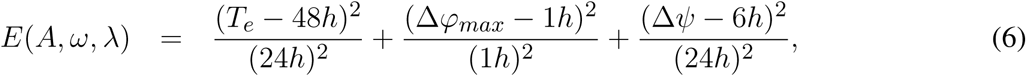

where *T*_*e*_, Δ*ϕ*_*max*_, Δ*ψ* represent half-time to re-entrainment, maximum phase-shift, and seasonal phase variability, respectively. The denominators can be regarded as tolerated ranges. If the values of *T*_*e*_, Δ*ϕ*_*max*_, and Δ*ψ* deviate 24h, 1h and 24*h* from their target values, a score of three results. All parameter sets discussed in this paper had optimized scores below 0.1, i.e., the constrains are well satisfied. Each quantity is defined and calculated as follows.

When the circadian oscillator is entrained to the *Zeitgeber* signal, their phase difference *ψ* = Ψ − *ϕ* (Ψ= 2*πt/T*: phase of the *Zeitgeber*) converges to a stable phase *ψ*_*e*_, that is called “phase of entrainment.” The half-time to re-entrainment *T*_*e*_ denotes the amount of time required for the oscillator to recover from a jet-lag. As the *Zeitgeber* phase is advanced by ΔΨ, the phase difference becomes *ψ* = *ψ*_*e*_ + ΔΨ. *T*_*e*_ quantifies how long it takes until the advanced phase is reduced to less than half of the original jet-lag (*i.e.*, |*ψ* − *ψ*_*e*_| < 0.5ΔΨ). In our computation, this quantity was averaged over 24 different times during the day, at which 6h-advanced jet-lag was applied. Next, the seasonal phase variability, which quantifies variability of the phase of entrainment over photoperiod from long day (16:8 LD) to short day (8:16 LD), is computed as Δ*ψ* =max_*κ*∈[1/3,2/3]_*ψ*_*e*_ − min_*κ*∈[1/3,2/3]_ *ψ*_*e*_. Finally, the maximum phase-shift is given by Δ*ϕ*_*max*_ = max_*ϕ*_ PRC(*ϕ*), where *PRC*(*ϕ*) represents phase response curve of the free-running oscillator, to which 6h light pulse is injected at its phase of *ϕ*.

To find optimal parameter values, the cost function was minimized by a particle swarm optimization algorithm (Eberhart and Kennedy, 1995; Trelea, 2003). Search ranges of the parameter values were set to *A*∈[0, 5], *ω*∈[2*π/*30, 2*π/*18], *λ*∈[0, 0.5]. Altogether 300 sets of parameter values were obtained. From the estimated parameters, the intrinsic period was obtained as *τ* = 2*π/ω*.

### 2.2 Simulations of jet-lag, phase response, and seasonality

Figure 1 illustrates our modeling approach. In Fig. 1a, the amplitude-phase equations (1) and (2) are visualized in the phase plane together with the driving *Zeitgeber* switching between 0 (dark) and 1 (light) for varying photoperiods. Two values of the amplitude relaxation rate *λ* illustrate how *λ* affects the decay of perturbations. Starting from an initial condition, small relaxation rate gives rise to a long transient until its convergence to limit cycle, while large relaxation rate exhibits only a short transient. Figure 1b shows the oscillations in a 3-dimensional phase space. Two coordinates (*x* and *y*) span the phase plane of the endogenous oscillator. The vertical axis represents the phase of the *Zeitgeber*. The red line marks the periodically forced limit cycle. The jump from 24 h to 0 h reflects simply the periodic nature of our daily time.

**Figure 1.**
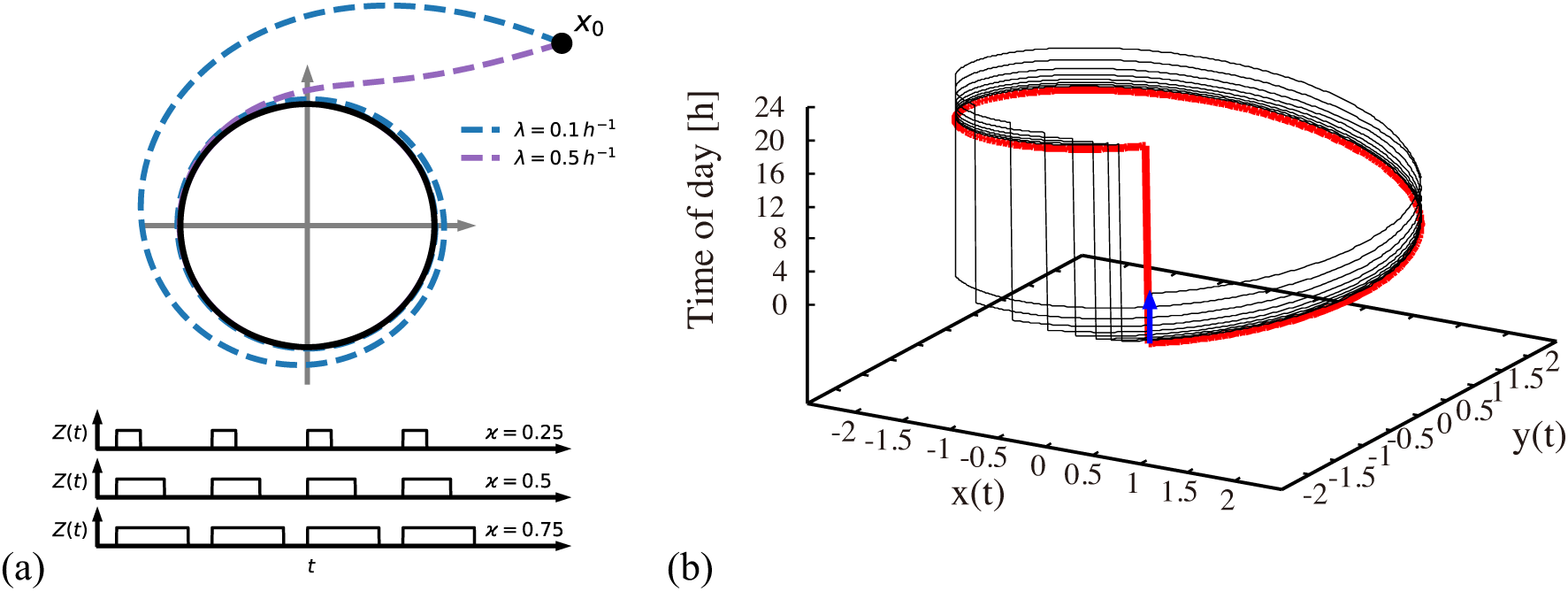
Visualization of the amplitude-phase model. (a) Schematic illustration of the amplitude-phase oscillator model forced by a *Zeitgeber* (light) signal of different photoperiods *κ*. Starting from the initial condition *x*_0_, a small relaxation rate (*λ* = 0.1 h^−1^) gives rise to long transient until its convergence back to the limit cycle, while transients for large relaxation rates (*λ* = 0.5 h^−1^) are short. (b) The re-entrainment process of the oscillator after its phase is shifted by a 6h-advanced jet-lag. The red line represents the jet-lag shifted trajectory that the system converges to. The blue arrow indicates the 6h jet-lag.

Interestingly, the relaxation after jet-lag (see also the more common presentations in Fig. 2) can be visualized as a transient convergence to the red limit cycle via the black line after a 6h phase change due to jet-lag (blue arrow). Such a relaxation might be accompanied by amplitude changes (not apparent) and by steady phase shifts from day to day (note that the jump from 24 h to 0 h is shifted day by day). After a few days, the red line is approached implying a vanishing jet-lag.

**Figure 2.**
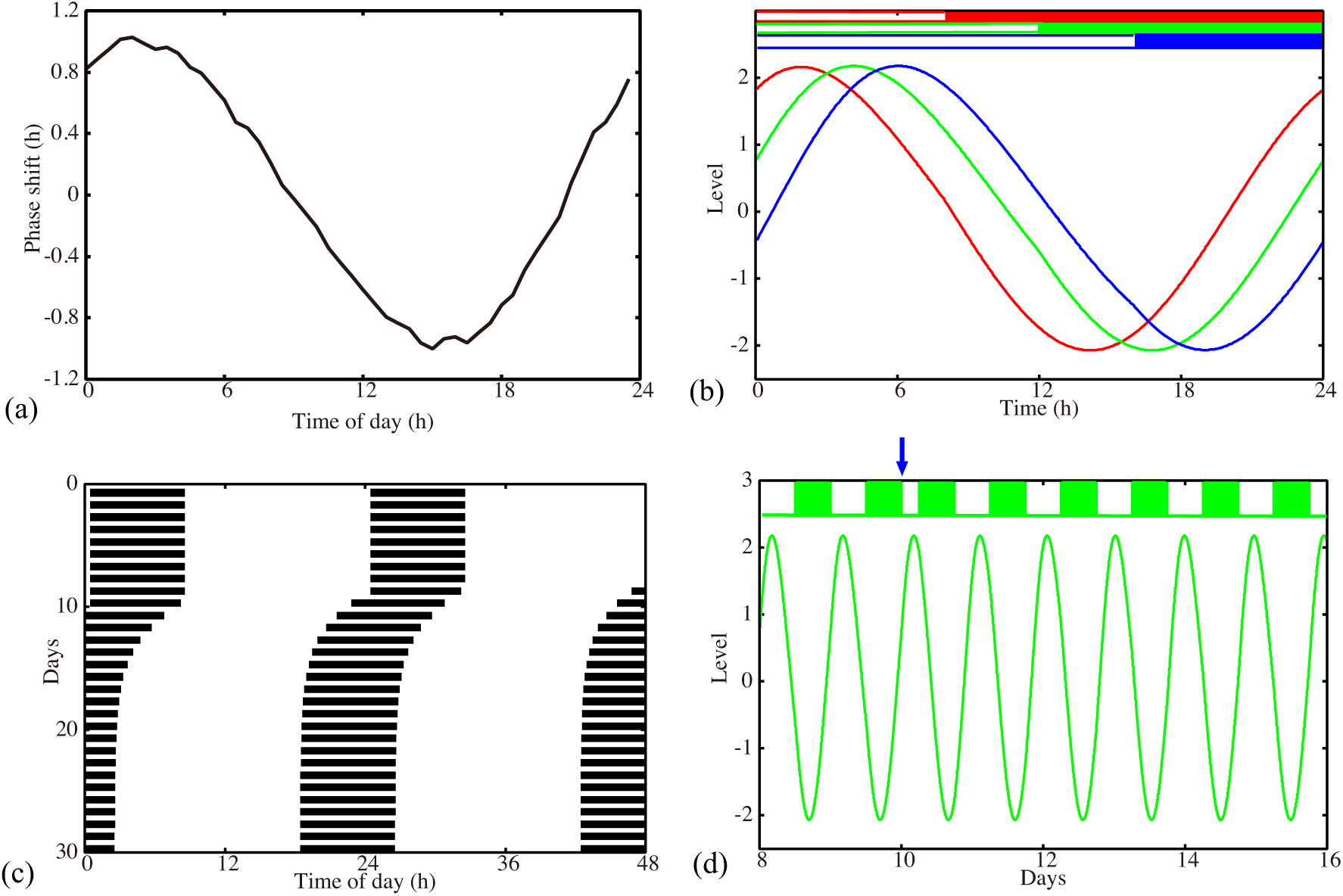
Properties of our amplitude-phase model for a representative optimized parameter set. (a) Phase response curve with respect to 6h light pulse. (b) Waveforms *x*(*t*) of the oscillator entrained to *Zeitgeber* signal with 8:16 LD (red), 12:12 LD (green), and 16:8 LD (blue). (c) Actogram drawn for the oscillators, to which a 6h advancing jet-lag was induced on day 10. (d) Time-trace *x*(*t*) of the oscillators, to which a 6h advancing jet-lag was induced on day 10. Model parameters: *τ* = 23.36h, *A* = 2.063, *λ* = 0.386h^−1^ with 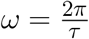.

## 3 RESULTS

### 3.1 Models reproduce small PRC, short jet-lag, and seasonality

We performed 200 successful parameter optimizations leading to an ensemble of parameter sets. We discuss in this section the parameters with a PRC of 1h delay and advance. In Figs. S1 & S2, we also present a parameter set obtained with a modified optimization: in that case, we requested a PRC with 2h delay and advance.

Figure 2 shows results for a representative model obtained via optimization. The PRC in Fig. 2a is almost sinusoidal with maximal delays and advances of 1h as requested by optimization. Simulations with different photoperiods are shown in Fig. 2b. It is evident that there are major phase shifts due to varying photoperiods. The jet-lag visualized in Fig. 2c is surprisingly short for such a relatively small PRC. Fig. 2d illustrates the re-entrainment after a jet-lag applied on day 10. Note, that no pronounced amplitude changes were observed.

It is remarkable that such simple models with just three free parameters can reproduce phenotypic features successfully. In particular, short jet-lag durations for quite small PRCs are surprising. In the following we exploit the ensembles of parameter sets to understand the underlying principles.

### 3.2 Optimization produced highly clustered parameter sets

In this section, we focus on the 200 parameter sets with the *±*1h PRCs exemplified in Fig. 2 (see Figs. S1 for *±*2h PRCs).

The possible search ranges for our parameters were quite large (periods between 18 and 30h, amplitudes between 0 and 5, and amplitude relaxation rates between 0 and 0.5*h*^−1^). The histograms from the optimized parameter sets demonstrate that the search lead to quite specific values: amplitudes of about 2.1*±*0.04, periods around 23.3*±*0.1 h, and large relaxation rates of 0.4*±*0.1 *h*^−1^.

The optimized amplitude can be easily understood from the constraint via PRCs: for a given pulse strength the PRC shrinks monotonically with increasing amplitude (Pittendrigh, 1981; Vitaterna et al., 2006). Consequently, the amplitude of about 2.1 implies a *±*1h PRC. Indeed, our optimizations with a *±*2h PRC lead to smaller amplitudes around 1.2.

Amplitude relaxation rates range between 0.1 and 0.5h^−1^. A value of 0.1h^−1^ corresponds to a half-life of the amplitude perturbations of about 7h, while a value of 0.5h^−1^ corresponds to a half-life of the amplitude perturbations of about 1.4h. Thus all values in the histogram imply relatively fast amplitude relaxation. In Abraham et al. (2010), we termed limit cycles with fast amplitude relaxation “rigid oscillators.” Interestingly, Comas et al. (2007) found that light pulses separated by 10h shift phases almost independently. This observation is consistent with fast amplitude relaxation rates. Jet-lag is a specific type of a transient (compare Fig. 1b). Thus it seems reasonable that fast amplitude relaxation helps to achieve short transients after a jet-lag.

The most surprising result of our optimization is the narrow range of intrinsic periods of about 23.3h. We provide in the discussion arguments that specific periods allow appropriate seasonal flexibility (compare Fig. 4). In short, at specific parts of *Arnold* onions (i.e. the entrainment regions in the *κ* − *T* parameter plane), the required 4h phase differences are found giving a reasonable phase shifts between 16:8 LD and 8:16 LD.

Figure 3d illustrates that the optimized parameter values are not independent. For example, shorter periods are associated with larger amplitude. A possible explanation is that short periods imply larger effective pulse strength (a 6h pulse is than relatively long) leading to larger amplitude in order to have the requested PRC amplitude.

**Figure 3.**
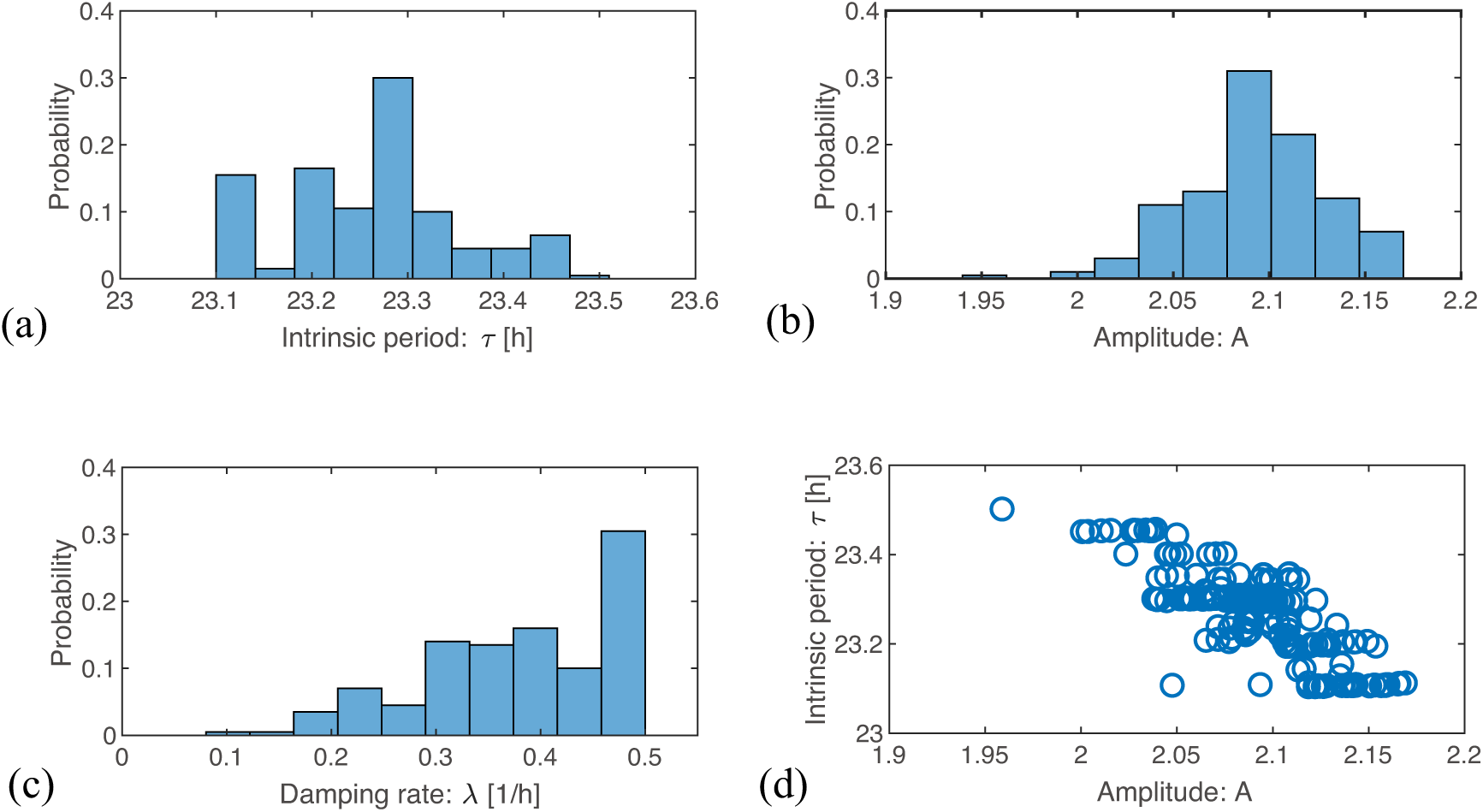
Parameter values in the optimized ensemble. (a-c) Distributions of the optimized parameter values for *τ* (= 2*π/ω*), *A*, and *λ*, respectively. (d) Scatter plots of amplitude *A* against intrinsic period *τ* drawn for the 200 sets of optimized parameters.

In order to evaluate the robustness of our optimization approach, we generated also 100 parameter sets with PRCs with about a 2h advance and delay. In these cases we found intrinsic periods of 24.6±0.1 h and amplitudes 1.15±0.1. The relaxation rates and amplitude-period correlations were similar to the results with PRCs of about 1h advance and delay, compare Fig. 3 and Fig. S1.

### 3.3 Arnold onions provide insights on the optimized parameters

To systematically investigate the impact of photoperiod (*κ*) and Zeitgeber period (*T*) on entrainment properties, we analyze in Figure 4 two *Arnold* onions for representative parameter sets with a short period and a *±*1 h PRC as well as a large period and a *±*2 h PRC. Interestingly, the *Arnold* onions are tilted, *i.e*. the DD periods at photoperiod *κ* =0 are smaller than the LL periods at photoperiod *κ* =1 as predicted by *Aschoff’s rule* for nocturnal animals (Aschoff, 1960). The largest entrainment range is found around a photoperiod of 0.5 as predicted by Wever (1964). As expected, a larger PRC implies a wider range of entrainment, compare sizes of the Arnold onion in Fig. 4 (a) and (b).

**Figure 4.**
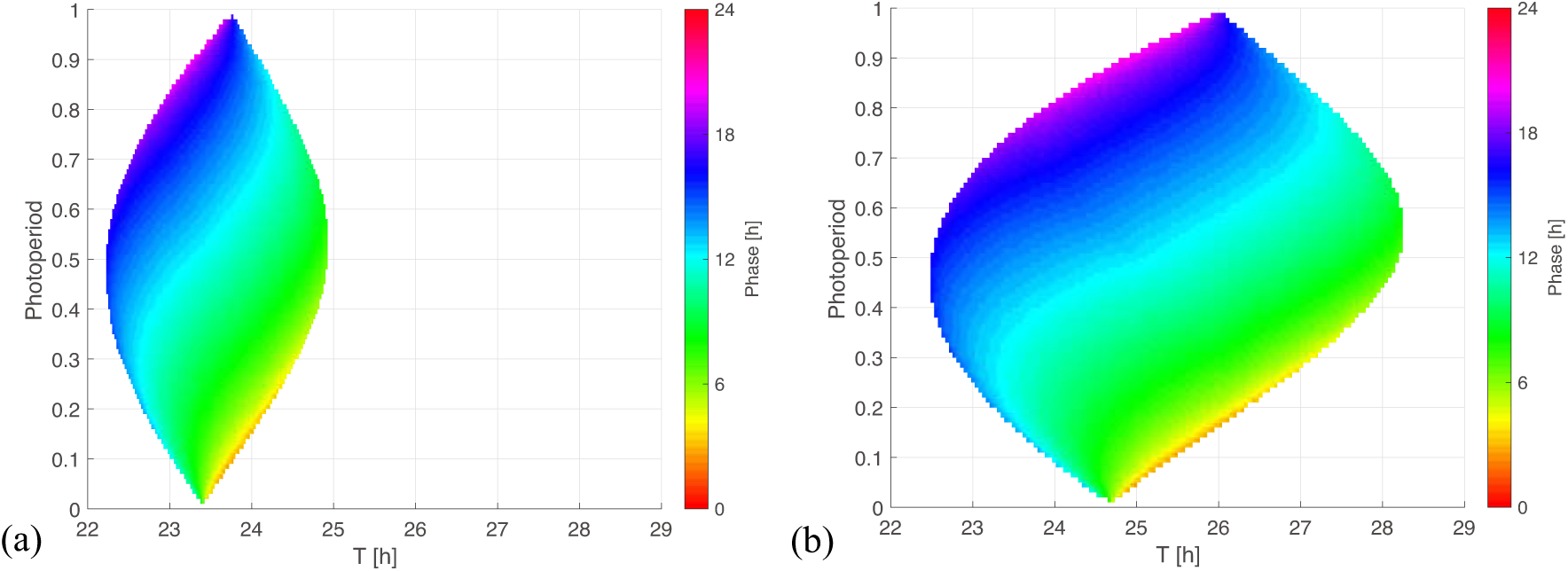
*Arnold* onions for the two PRC constraints. (a,b) 1:1 entrainment ranges in the *κ-T* parameter plane (*Arnold* onions). Entrainment phases were determined by numerical simulations and have been color-coded within the region of entrainment. Panel (a) depicts an *Arnold* onion for an optimized parameter se with a ±1h-PRC, a short-period *τ* = 23.36h, *A* = 2.063, and *λ* = 0.386h^−1^. Panel (b) shows an *Arnold* onion for an optimized parameter set with a ±2h-PRC, a long-period *τ* = 24.64h, *A* = 1.144, *λ* = 0.50h^−1^.

The phase of entrainment is color-coded in Fig. 4. It is evident that the phases vary strongly with the external period *T*. There are theoretical predictions that phases change by about 12h (Wever, 1964; Granada et al., 2013; Bordyugov et al., 2015) in the horizontal direction. Interestingly, the variation of photoperiods in the vertical direction implies also very pronounced variations of the entrainment phase. Consequently, we could find many parameter sets with about 4h phase shift between photoperiods of *κ* = 1/3 (8:16 LD) and of *κ* = 2/3 (16:8 LD).

## 4 DISCUSSION

### 4.1 Optimized models reproduce phenotypic features

The circadian clock of vertebrates is characterized by a relatively small type 1 PRC, by a narrow entrainment range, by fast recovery from jet-lag, and by pronounced seasonal flexibility. We addressed the question whether or not these phenotypic features can be reproduced by simple amplitude-phase models with just 3 free parameters: period, amplitude, and relaxation time. To our surprise, we found many parameter sets via global optimization that reproduce the phenotypic features.

The availability of many parameter sets derived from random optimization allows extraction of essential properties of successful models. It turned out that the amplitude *A* is adjusted to reproduce small PRCs for a given *Zeitgeber* strength *Z*. According to limit cycle theory (Pavlidis, 1969; Peterson, 1980), the strength of a perturbation is governed by the ratio *Z/A* of *Zeitgeber* strength to amplitude. This implies that large limit cycles exhibit small PRCs for a fixed *Zeitgeber* strength (Vitaterna et al., 2006).

In all suitable models, we found relatively fast amplitude relaxation rates with half-lives of perturbations below 5h. This “rigidity” of limit cycles (Abraham et al., 2010) can support fast relaxation to the new phase after jet-lag (compare Fig. 1b). Interestingly, Comas et al. (2007) found that light pulses separated by 10h act almost independently. This observation is consistent with fast relaxation rates after light pulses.

In order to reproduce seasonality we optimized our model under the constraint that 16:8 and 8:16 LD cycles have entrainment phases that are about 4h apart. This implies that the phase could follow dusk or dawn (Daan and Aschoff, 1975). In other words, we requested that the entrainment phase depends strongly on the photoperiod. As illustrated in Fig. 4, such a strong dependency is indeed reproduced by our simple amplitude-phase models. Our optimization procedure selected specific periods that lead to a 4h phase variation between photoperiods *κ* = 1/3 and *κ* = 2/3. Note, that other periods can give large phase differences as well (compare the large vertical phase variations in Fig. 4).

### 4.2 Relevance of phenomenological amplitude-phase models

Simplistic models as studied in this paper are quite generally applicable. In principle, they could be used to describe single cells, tissue clocks, and organismic data. For single cells, damped stochastic oscillators can represent the observations also surprisingly well (Westermark et al., 2009). Such models have vanishingly small amplitudes, smaller relaxation rates, and are driven by stochastic terms. Otherwise their complexity is comparable to our models discussed above.

Complex models with multiple gene-regulatory feedback loops (Mirsky et al., 2009; Pokhilko et al., 2010; Relógio et al., 2011; Woller et al., 2016) could be reduced to amplitude-phase models simply by extracting periods, amplitudes, and relaxation rates from simulations. However, in such cases the amplitudes are not uniquely defined, since there are many dynamic variables.

### 4.3 How to define circadian amplitudes?

This difficulty to define amplitudes points to a general problem in chronobiology. Most studies focus on periods and entrainment phases. Limit cycle theory emphasizes that amplitudes are essential to understand PRCs and entrainment. It is, however, not evident which amplitudes properly represent the limit cycle oscillator. Some studies consider gene expression levels (Lakin-Thomas et al., 1991; Wang et al., 2019) or reporter amplitudes (Leise et al., 2012), and other studies quantify activity rhythms (Bode et al., 2011; Erzberger et al., 2013). Since the ratio of *Zeitgeber* strength to amplitude *Z/A* governs PRCs and entrainment phases, we suggest that amplitudes could be quantified indirectly: the stronger the response to physiological perturbations, the smaller the amplitude. This approach leads to the concept of strong and weak oscillators (Abraham et al., 2010; Granada et al., 2013). Strong oscillators are robust and have small PRCs and entrainment ranges but large phase variability (Granada et al., 2013). In this sense, wild-type vertebrate clocks represent strong oscillators in contrast to single cell organisms or plants. Indeed, the review of Aschoff and Pohl (1978) demonstrates impressively these properties.

Interestingly, a reduction in relative amplitudes (i.e., amplitudes as a fraction of the mesor) can reduce jet-lag drastically, since resetting signals are much more efficient (Yamaguchi et al., 2013; An et al., 2013; Jagannath et al., 2013).

### 4.4 Arnold onions quantify entrainment

As shown in Fig. 4, *Arnold* onions represent in a compact manner entrainment ranges and phases of entrainment. Astonishingly, even quite basic models lead to really complex variations of entrainment phases. As expected, the period mismatch *T* –*τ* has a rather strong effect on the entrainment phase. This reflects the well-known feature that short intrinsic periods *τ* have earlier entrainment phases (“chronotypes”) (Pittendrigh and Daan, 1976a; Merrow et al., 1999; Duffy et al., 2001). These associations are reflected in the horizontal phase variations in the *Arnold* onion. Interestingly, the vertical phase variability is also quite large. This observation demonstrates that also the effective *Zeitgeber* strength *Z/A* and the photoperiod affect the phase of entrainment strongly. Consequently, the expected correlations between periods and entrainment phase could be masked by varying amplitudes, *Zeitgeber* strength, and photoperiods. In other words, chronotypes are governed by periods only if relative *Zeitgeber* strength and photoperiod are kept constant.

The complexity of entrainment phase regulation indicates that generic properties of coupled oscillators can provide useful insight in chronobiology. In particular, basic amplitude-phase models can help to understand the control of jet-lag and seasonality.

## AUTHOR CONTRIBUTIONS

H. H. designed the study. I. T. T. performed the model simulations. I. T. T., B. A., C. S., and H. H. discussed the results. H. H. wrote the main text. I. T. T., B. A., and C. S. revised the text.

## ACKNOWLEDGMENTS

The authors thank Serge Daan (*†*), Achim Kramer, and Adrian Granada for stimulating discussions.

---

**Figure S1.**
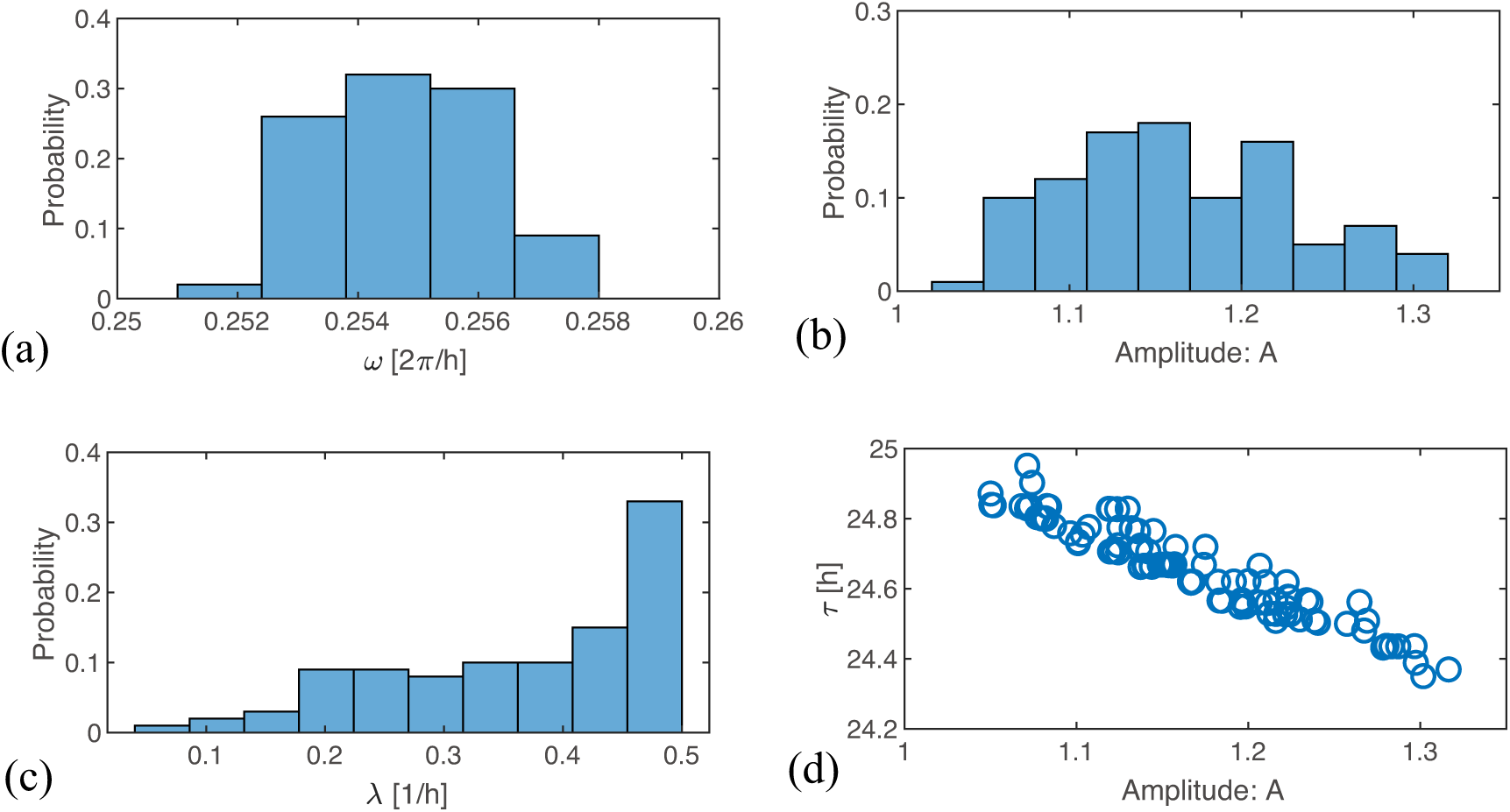
Results of parameter optimization based on cost function 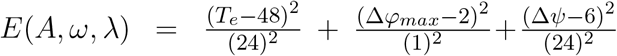, where *±*2h PRC was requested. (a-c) Distributions of the 100 optimized parameter sets for *τ* (24.6±0.16 h), *A* (1.2±0.07), *λ* (0.37±0.12), respectively. (d) Scatter plots of amplitude *A* against intrinsic period *τ* drawn for 100 sets of optimized parameters.

**Figure S2.**
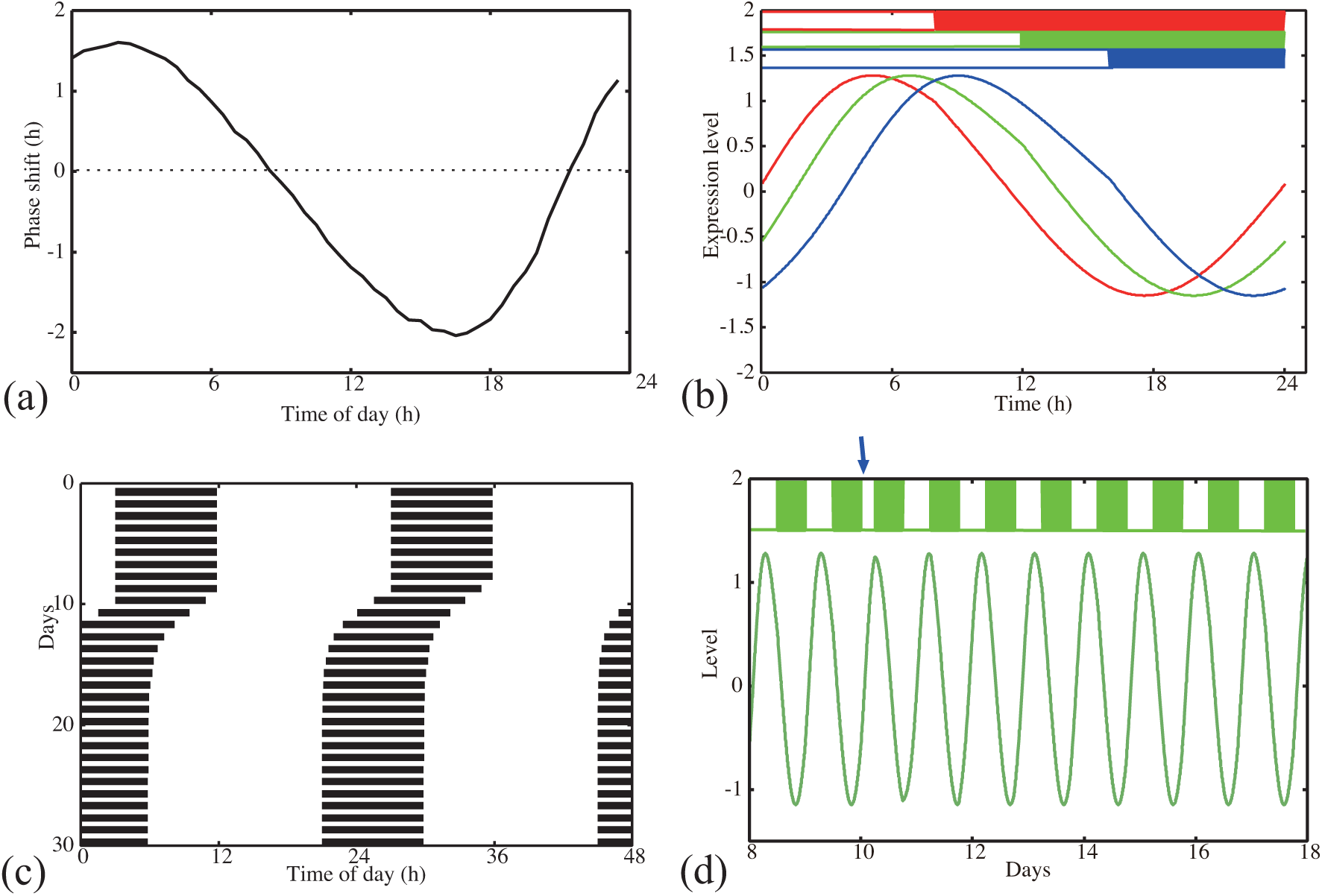
Simulation of the amplitude-phase oscillator model using one of the 100 parameter sets optimized for a ±2h PRC. (a) Phase response curve with respect to 6h light pulse. (b) Waveforms *x*(*t*) of the oscillator entrained to *Zeitgeber* signal with 8:16 LD (red), 12:12 LD (green), and 16:8 LD (blue). (c) Actogram drawn for the oscillator, to which a 6h advancing jet-lag was induced on day 10. (d) Time-trace *x*(*t*) of the oscillators, to which a 6h advancing jet-lag was induced on the 10th day. Parameter values: *τ* = 24.64h, *A* = 1.144, *λ* = 0.50h^−1^.

